# CGIAR BARLEY BREEDING TOOLBOX: A diversity panel to facilitate breeding and genomic research in the Developing World

**DOI:** 10.1101/2022.08.13.502796

**Authors:** O. Bouhlal, A. Visioni, R.P.S. Verma, M. Kandil, S. Gyawali, F. Capettini, M. Sanchez-Garcia

## Abstract

Despite the reduced cost of the new genotyping technologies, many public and private breeding programs, mostly in developing countries, still cannot afford them, hindering their research. The objective of the present study was to identify and assemble an Association Mapping panel of widely diverse barley genotypes to serve as a barley breeding toolbox, especially for the Developing World. The main criteria were: i) to be representative of the germplasm grown in the Developing World; ii) to cover a wide range of genetic variability and iii) to be of public domain. For it, we assembled and genotyped a Global Barley Panel (GBP) of 530 genotypes representing a wide range of row-types, end-uses, growth habits, geographical origins and environmental conditions. The GBP accessions were genotyped using the barley Infinium iSelect 50K chip. A total of 40,342 SNP markers were polymorphic and displayed an average polymorphism information content (PIC) of 0.35, with 66% of them exceeding PIC=0.25. The analysis of the population structure identified 8 sub-populations mostly linked to the geographical origin of the lines (Europe, Australia, USA, Canada, Africa, Latin-America, ICARDA and others), four of them of significant ICARDA origin. The 16 allele combinations at 4 major flowering genes (HvVRN-H3, HvPPD-H1, HvVRN-H1 and HvCEN) explained 11.07% genetic variation and were linked to the geographic origins of the lines studied. Among origins, ICARDA material (n= 257) showed wide diversity as revealed by the highest number of polymorphic loci (99.76% of all polymorphic SNPs in GBP), number of private alleles and Nei’s gene diversity and the fact that ICARDA lines were present in all 8 sub-populations and carried all 16 allelic combinations at phenology genes. Due to their genetic diversity and their representativity of the germplasm adapted to the Developing World, 312 ICARDA lines and cultivated landraces were pre-selected to form the CGIAR Barley Breeding Toolbox (CBBT). Using the genotypic data and the Mean of Transformed Kinships method, we assembled the CBBT, an Association Mapping panel of 250 accessions capturing most of the allelic diversity in the global panel. The CBBT preserves a good balance between row types as well as a good representation of both allelic combinations identified at most important phenological loci and sub-populations of the GBP. The CBBT lines together with their genotypic data will be made available to breeders and researchers worldwide to serve as a collaborative tool to underpin the genetic mechanisms of traits of interest for barley cultivation.

## 1. Introduction

The new advances in genotyping technologies and genomic research applied to breeding (Genome Wide Association Studies and more recently Genomic Selection) have the potential to bring the largest yield productivity increase and stability improvement since the Green Revolution. The efficient use of these new tools is important to attain the production needed to feed the increasing population in a scenario of climate instability. These new approaches have been widely adopted thanks to new low-cost and high throughput genotyping technologies (Paux et al., 2010; Rimbert et al., 2018). However, despite the reduced cost, many public and private breeding programs, mostly in developing countries, still cannot afford them and as a result, a new technological gap between the developing and developed world has arisen.

Barley (*Hordeum vulgare* L.) covers 50Mha worldwide, ca. 20Mha of them in developing countries (FAOSTAT 2022). Its drought tolerance and integration in the traditional crop-livestock farming systems make this crop an integral part of the strategies to cope with Climate Change, especially in the Dry Areas (Sanchez-Garcia 2021). With the mandate to develop new barley varieties and cutting-edge research for the developing World, the CGIAR Global Barley Breeding program (ICARDA), in collaboration with national partners, has successfully developed, since 1977, more than 250 cultivars released and adopted by farmers. Genomic studies have been an integral part of the research carried out in the program to underpin the genetic control of traits of interest such as drought tolerance (Varshney et al., 2012), disease resistance (Visioni et al., 2018, 2020) and nutritional and malting quality (Gyawali et al., 2017; Boulal et al. 2021) among others. This research has been integrated into the breeding program and the new germplasm developed carries novel QTL for traits of interest, benefiting partners Worldwide. However, despite the large network of testing locations and phenotyping capacities that the program uses - mostly in Morocco, Egypt, Ethiopia, Lebanon and India - the capacity to identify new QTL is still limited, especially for stresses not occurring at the testing locations. A collaborative network of scientists, particularly from the developing world, testing the same germplasm across local conditions and stresses and for traits of interest could help identifying new relevant alleles that will then be available to the whole network.

Several highly diverse barley populations have been assembled in the last 30 years to study the genetic control of traits. Most of these populations consisted in core collections of major genebanks (Van Hintum et al., 1995; Igartua et al., 1998; Muñoz-Amatriaín et al., 2014) or even multilateral efforts such as the International Barley Core Collection in 1989 (Knüpffer and van Hintum, 2003). Also, collections of modern varieties and elite germplasm have been used (Looseley et al., 2020), however, many of these populations are not representative of the barley varieties adapted to the developing world or are too large to be grown with limited resources. In addition, only a limited number of them take advantage of the new genotyping technologies that allow the use of large number of representative stable markers such as the 50k Illumina Infinium iSelect genotyping array for barley (Bayer et al., 2017) or are not freely available to the international community.

When studying subpopulation genetic differentiation, Muñoz-Amatriaín et al. (2014), found that many differentially selected genomic regions are coincidental with, or near to known loci involved in flowering time and spike row number. Russell et al. (2016) reported similar differentiation between two and six-row germplasm close to the location of *VRS1* on chromosome 2H (Komatsuda et al., 2007) and INTERMEDIUM-C on chromosome 4H (Ramsay et al., 2011) known as controlling the number of fertile florets in barley. In fact, the two spike morphologies have traditionally been separated geographically, and this separation has been reinforced by modern plant breeding (Muñoz-Amatriaín et al., 2014). For it, and building on the germplasm exchange network of international collaborators; the consistent use of genebank material in the breeding program and recent efforts to assemble relevant association panels (Amezrou et al., 2018; Verma et al., 2021), the Global Barley Breeding Program has developed and genotyped a global panel of 530 barley genotypes. This population includes the different barley uses (feed, forage, malt and food), morphophysiological traits and phenology patterns to provide meaningful diversity to collaborators irrespective of their target environments, local preferences and end-use breeding targets.

Similarly to recent panels assembled for other cereals, such as durum (Mazzucotelli et al., 2020), we aim to take advantage of the extensive expertise of ICARDA on germplasm exchange to make available to the international community an Association Mapping panel of 250 highly diverse barley genotypes. The main objective of the present study is to identify and assemble an Association Mapping panel of 250 barley genotypes that represent the global diversity to serve as a barley breeding toolbox, especially for the Developing World. For it, the main criteria to select among the lines in the Global panel was: i) to be representative of the germplasm grown in the Developing World; ii) to cover a wide range of genetic variability, morphophysiological traits and phenology groups and iii) be of public domain or international public goods to facilitate germplasm exchange.

## 2. Materials and Methods

### 2.1. Plant material

A Global Barley Panel (GBP) of 530 genotypes including elite lines, cultivars, and landraces from different geographic origins was assembled. The GBP includes a significant number of elite lines and cultivars from the ICARDA Global Barley Breeding Program of the CGIAR, both from the ICARDA program based in Syria, now in Morocco and Lebanon, and from the ICARDA/CIMMYT Latin-America program based in Mexico, merged during the first decade of the 2000s. The set represents a worldwide spectrum of barley genetic variance with diversity of row-types, end-uses, growth habits and geographical and environmental distribution. Out of the 530 genotypes (292 two-row and 238 six-row types), 288 are from the ICARDA Global Barley Breeding Program, 61 from the United States of America (USA), 48 from Europe, 40 from India, 23 from Australia, 19 from Canada, 12 from Africa, 6 from Latin America and 16 from other countries across the globe. The Panel also includes 17 landraces from different countries (**Figure 1**). A detailed list of the genotypes used is provided in **Supplementary Table S1**.

**Figure 1.**
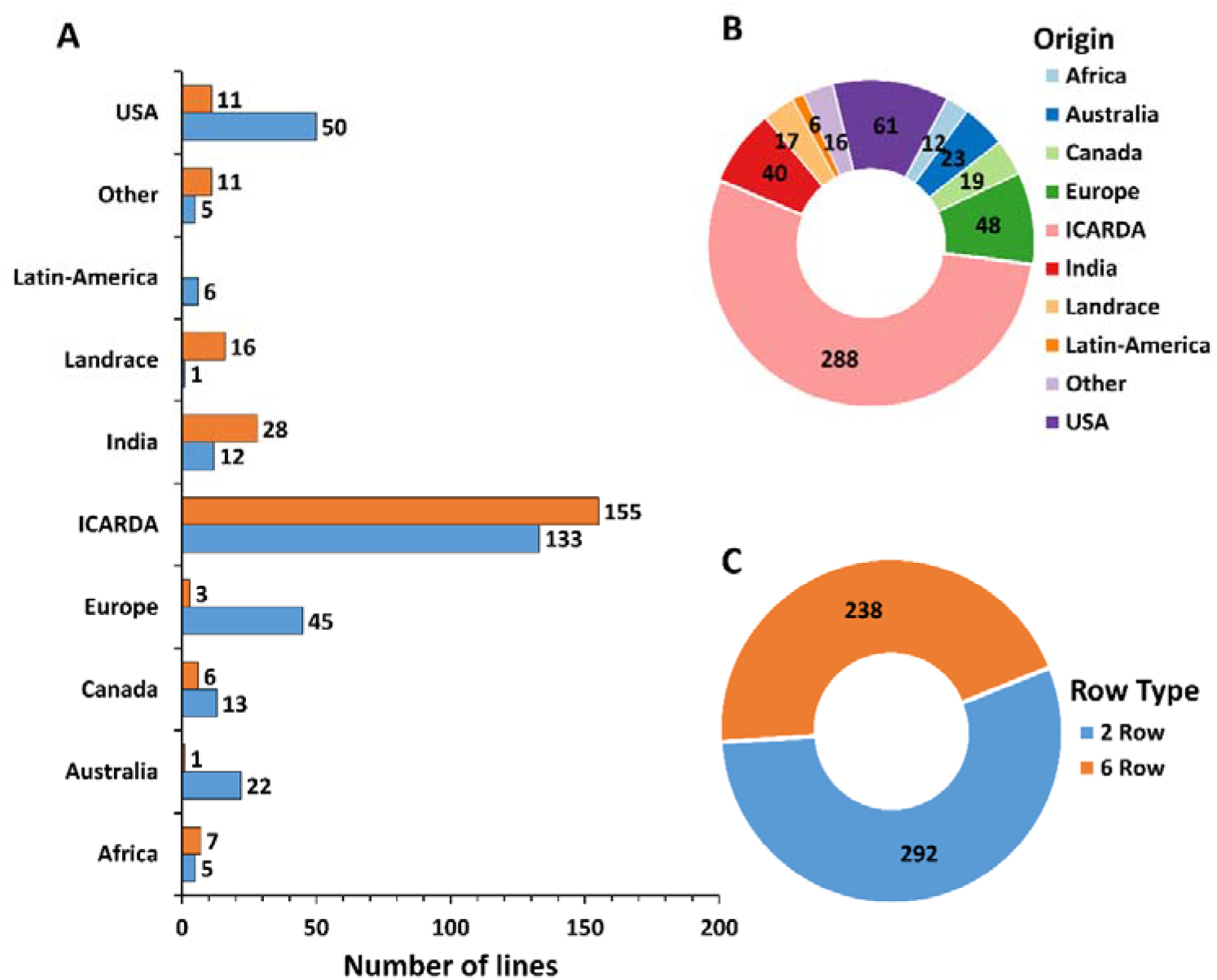
Characterization of the 530 genotypes used based on row number (A and C) and geographic origin (A and B).

### 2.2. DNA Extraction and SNP Genotyping

The genotyping work was done in two events. The first 266 entries were genotyped as described in Verma et al. (2021). For the second event, genomic DNA was isolated from lyophilized 2-week-old leaf tissue from a single plant from each genotype as described in (Slotta et al., 2008). The GBP was genotyped using the recently developed barley Infinium iSelect 50K chip (Illumina, San Diego, California, USA) (Bayer et al., 2017) by TraitGenetics GmbH using the manufacturer’s guidelines. Out of 43,461 scorable SNP markers (Bayer et al., 2017), 40,342 were polymorphic (**Supplementary Table S2**). A final set of 36,253 SNPs was obtained after removing markers with minor allele frequencies (MAF) < 5% and markers with > 10% of missing data. The distribution of the filtered SNPs across the seven barley chromosomes (**Figure 2A)** was illustrated using the *CMplot* package (Yin et al., 2021) for R statistical software (R Core Team, 2019).

**Figure 2.**
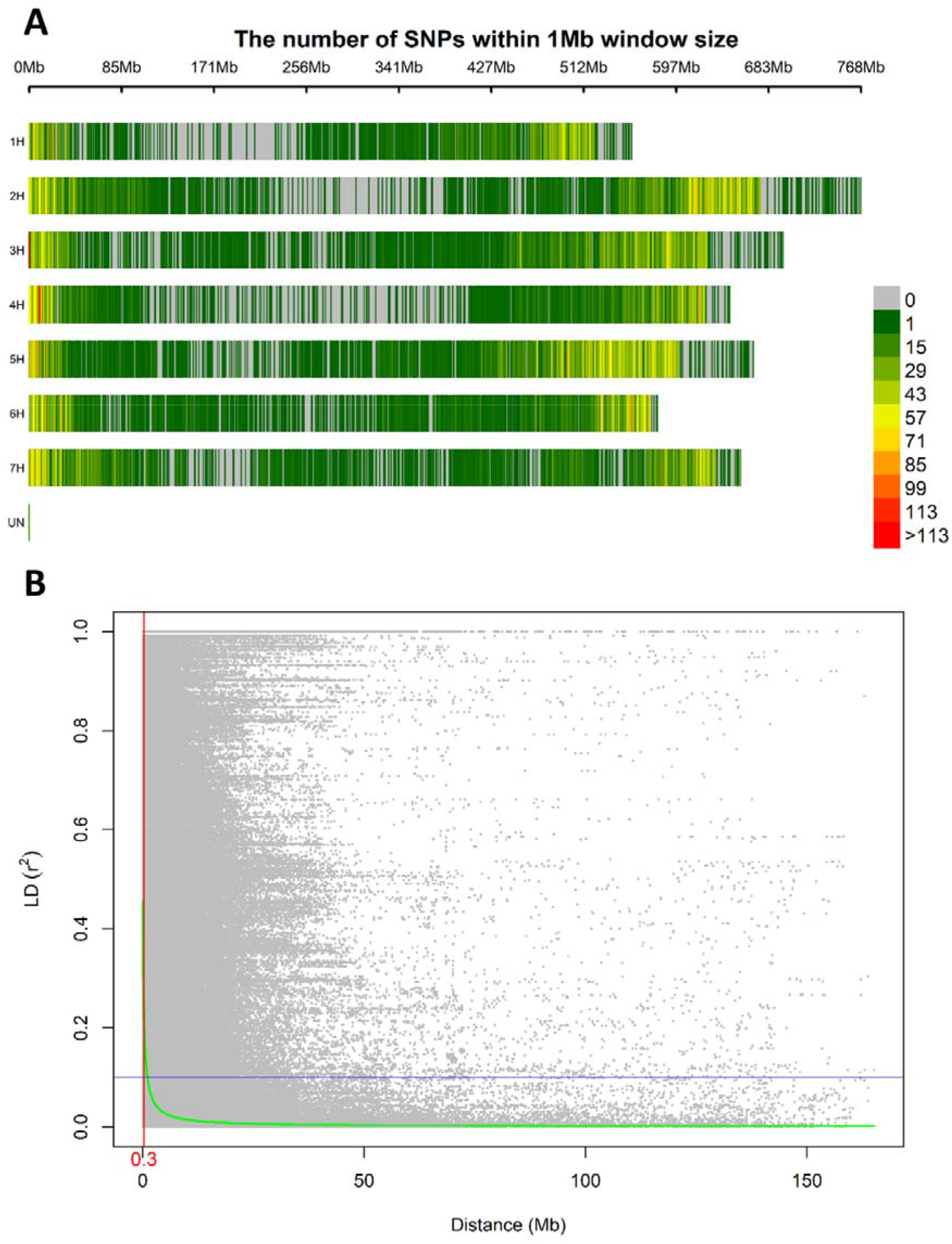
Distribution in the barley genome of 40,342 SNPs markers based on the physical map (A). Linkage disequilibrium expressed per distance between markers in the same chromosome (B). The linkage decay distance is shown in red.

### 2.3. Linkage Disequilibrium Analysis

From the set of 40,342 SNP markers, a subset of 40,256 polymorphic markers with known positions was selected to perform the linkage disequilibrium (LD) analysis for each chromosome using TASSEL 5.0 software (Bradbury et al., 2007) and a sliding window of 50 SNPs. The LD for locus pairs within the same chromosome was estimated as the squared allele frequency correlations estimates (*R*^2^) and significant corresponding p-values ≤ 0.01. The R Statistical Software (R Core Team, 2019) was used to estimate the extent of LD by non-linear regression analysis based on all intrachromosomal *R*^2^ values as described in Hill and Weir (1988) and implemented in Remington et al. (2001). The distribution and extent of LD were visualized by plotting the *R*^2^ values against the genetic distance between same-chromosome markers (Mb).

### 2.4. Genetic Diversity and Population Differentiation

To estimate the genetic diversity within the GBP, a Principal Component Analysis (PCA) was conducted with the filtered SNP maker set using TASSEL version 5.0 (Bradbury et al., 2007), and the first two eigenvectors were considered. The unrooted neighbor-joining (NJ) clustering algorithm under the Provesti’s absolute genetic distances was applied using the R package *poppr* (Kamvar et al., 2015) to investigate the relationship among barley accessions. The R packages, *ggtree* (Yu, 2020) was used to visualize the NJ phylogenetic tree. Analysis of molecular variance (AMOVA) implemented in the R package *poppr* (Kamvar et al., 2015) was used to assess the genetic diversity between and within major clusters (row types and geographic origins) defined by PCA and the NJ clustering. For AMOVA of geographical origin, we excluded the “Other” group since it includes cultivars from different origins. The population differentiation was assessed based on population pairwise Fst calculated using *hierfstat* package (Goudet, 2005). Diversity indices analyses including Nei’s unbiased gene diversity (Nei, 1978) and allelic richness were calculated using the *“locus_table”* and “*poppr*” function of *poppr* R package (Kamvar et al., 2014).

### 2.5. Diversity analysis at flowering loci

AMOVA was used to assess the extent of genetic diversity existing within the 530 barley genotypes based on flowering genes. For it, SNP markers associated with major flowering time genes in barley were selected: BK_05 (T/C) associated to *Hv*VRN-H3 (FT1) gene, BK_14 (A/G) associated with *Hv*PPD-H1 gene, BK_17 (C/G) associated to *Hv*VRN-H1 gene, and BOPA2_12_30265 (A/G) associated with *Hv*CEN gene (Comadran et al., 2012). AMOVA tests were used to calculate the variation explained by each of the SNP markers associated with flowering genes and the 16 allelic combinations (AC) of the 4 markers. The list and detailed information of the selected SNPs are provided in **Supplementary Table S4.** AMOVA was conducted using the R package *poppr* (Kamvar et al., 2014). The AC obtained from the SNPs were used to identify germplasm classes based on their earliness and lateness. LD parameter (*R*^2^) calculated by TASSEL software (Bradbury et al., 2007) was used to test whether the selected SNPs were in LD with each other (**Supplementary Table S3**).

### 2.6. Population Structure

The 36,253 filtered SNP markers were used to calculate individual admixture coefficients using the sparse Non-negative Matrix Factorization (sNMF) algorithm implemented in the R package *LEA* (Frichot and François, 2015). This method was specifically developed to estimate individual admixture coefficients on large genomic datasets. The sNMF algorithm estimates ancestry independently for each individual and does not require prior assumptions about population membership. We used the cross-entropy criterion from the *snmf* function (Alexander and Lange, 2011; Frichot et al., 2014) to test several putative populations (K) ranging from K=2 to K=15. For each K we set the number of runs to 10, alpha to 10, tolerance to 10^-5^, and the iterations number to 200. Lines with strong admixture were defined as those showing less than 70% of identity (membership) with any ancestry in the model (**Supplementary Table S5**).

### 2.7. Genetic sub-setting

To select a subset that represents the diversity of the population and at the same time provides an optimized set of entries for association mapping studies we used the *R* package *GeneticSubsetter* (Beukelaer et al., 2012; Graebner et al., 2016). For it, we used the *Local Search* subsetting option that produces single-genotype replacements repeatedly to multiple random starting subsets, until no more single-genotype replacements can be made with 100 iterations. The subsetting criteria used was Mean of Transformed Kinships (MTK) which measures the kinship of genotypes in a subset and identifies the most dissimilar set of genotypes, a favorable trait for GWAS studies.

## 3. Results

### 3.1. SNP density and linkage disequilibrium

A total of 40,342 polymorphic SNP markers were retained after removing monomorphic markers and used for further genetic analysis. Only 86 SNP lacked chromosomal and physical map position information. The highest number of markers per chromosome was observed in chromosome 5H followed by 2H with 7,545 and 6,718 SNP respectively while chromosome 1H had the lowest number of SNPs amongst the seven chromosomes with 4,415 SNP. The MAF ranged from 0.26 to 0.28 with chromosome 4H having the lowest average MAF. The PIC values ranged from 0.34 to 0.36 among chromosomes and an average of 8.54 SNPs per Mbp was calculated (**Table 1**). Diversity statistics computed for each SNP are summarized in (**Supplementary Table S3**). The minimum PIC value for SNPs was 0.003 with an average of 0.35. Most of the markers (66.12%) displayed PIC values exceeding 0.25, indicating the informativeness of the genotyping in our population. The SNPs density along the seven barley chromosomes was higher in proximal and distal portions compared to pericentromeric regions (**Figure 2A**). The mean R^2^ values for the whole genome decreased with increasing pairwise distance. In general, LD showed a fast rate of decline, and the decay distance was approximately 0.3 Mbp (**Figure 2B**).

**Table 1.**
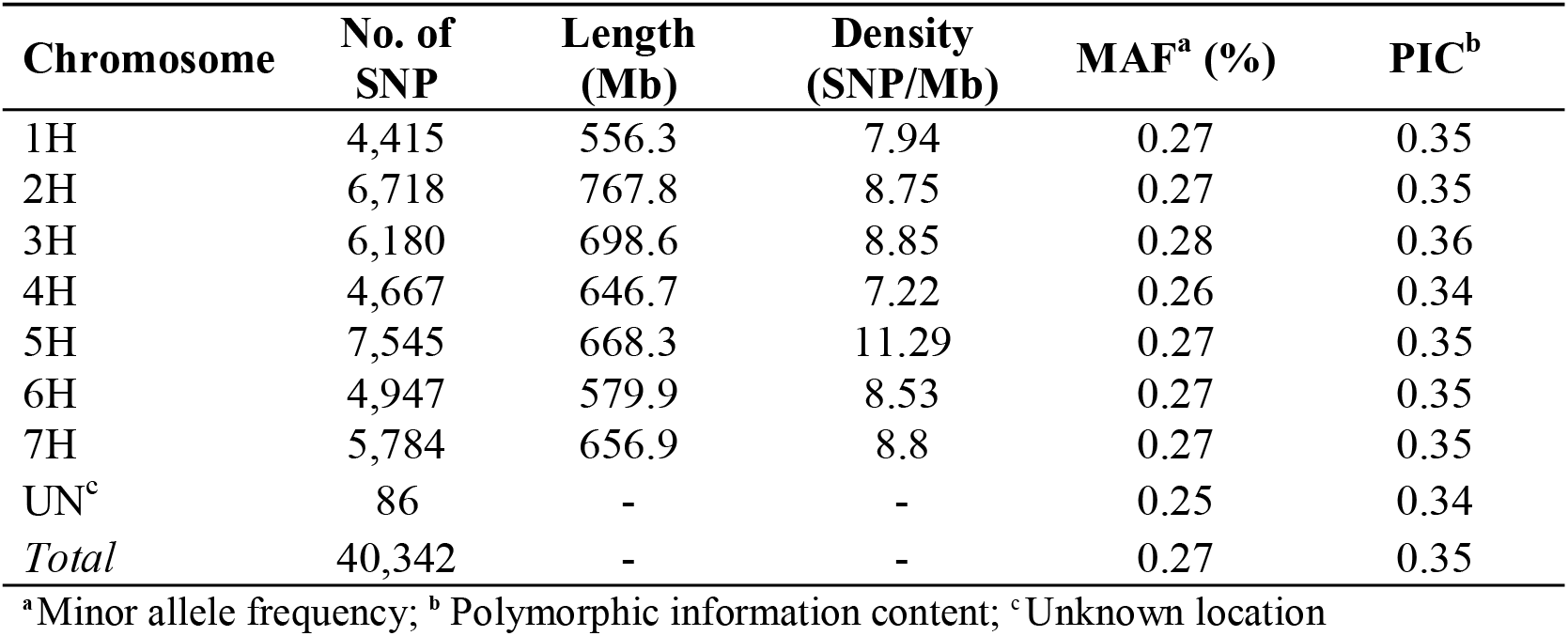
Genome coverage of a 40,342 SNP marker dataset used expressed for each chromosome and genome distribution across the seven barley chromosomes.

| Chromosome | No. of SNP | Length (Mb) | Density (SNP/Mb) | MAF<sup>a</sup> (%) | PIC<sup>b</sup> |
| --- | --- | --- | --- | --- | --- |
| 1H | 4,415 | 556.3 | 7.94 | 0.27 | 0.35 |
| 2H | 6,718 | 767.8 | 8.75 | 0.27 | 0.35 |
| 3H | 6,180 | 698.6 | 8.85 | 0.28 | 0.36 |
| 4H | 4,667 | 646.7 | 7.22 | 0.26 | 0.34 |
| 5H | 7,545 | 668.3 | 11.29 | 0.27 | 0.35 |
| 6H | 4,947 | 579.9 | 8.53 | 0.27 | 0.35 |
| 7H | 5,784 | 656.9 | 8.8 | 0.27 | 0.35 |
| UN<sup>c</sup> | 86 | - | - | 0.25 | 0.34 |
| Total | 40,342 | - | - | 0.27 | 0.35 |

<sup>a</sup>Minor allele frequency; <sup>b</sup> Polymorphic information content; <sup>c</sup> Unknown location

### 3.2. Genetic Diversity and population differentiation

We conducted PCA using the filtered set of 36,253 SNP markers to evaluate the genetic diversity of the 530 barley genotypes. The PCA of the GBP showed a clear differentiation based on row type and geographic origin. The first axis of the PCA, explaining 8.6% of the genotypic variance, separated the genotypes according to their row type, being the 2-row genotypes located towards the negative side of the axis and 6-row ones in the positive (**Figure 3A**). According to the phylogenetic tree (**Figure 3C**), two-rowed genotypes were grouped into five subgroups, the first was composed mainly of entries from Australia and ICARDA; the second was mostly from USA; the third grouped genotypes from Canada, USA and ICARDA; the fourth was predominantly from Europe, and the subgroup 7 was composed mainly of ICARDA entries. The six-rowed barley genotypes were split into three groups, with the first one (subgroup 5) composed of ICARDA, India, North and South America, the second (subgroup 6) and third (subgroup 8) represented two genetically different subgroups of ICARDA genotypes, as observed in the two-rowed genotypes (**Figure 3B**).

**Figure 3.**
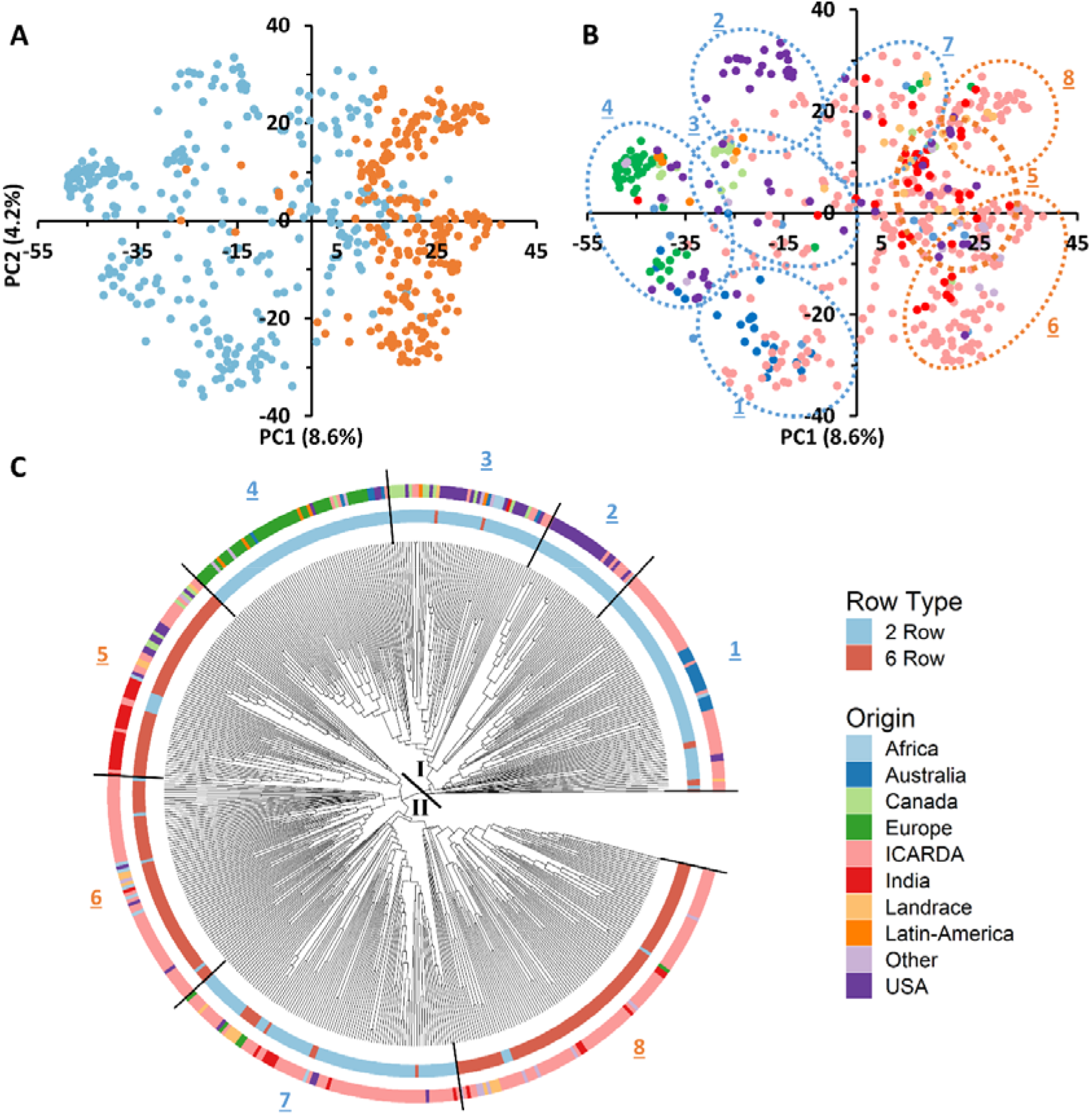
Genotypic and geographic diversity of the global barley panel used. Principal component analysis (PCA) of SNP markers displays the spatial distribution of the 530 genotypes (A, B). Individuals are shown with colored round symbols based on their row type (A) and geographic origin (B). The phylogenetic tree of the 530 barley genotypes based on 36,253 SNPs (C). The 530 accessions are colored according to their row type (inner circle) and geographic origin (outer circle).

The genetic variation between and within groups was further explored through AMOVA tests and the analysis of genetic diversity parameters (**Table 2** and **Table 3**, respectively). AMOVA revealed that the row type grouping accounted for more than 10.6 % of the whole genetic variation (**Table 2**). This result supports the grouping defined by the first PC in the PCA (**Figure 3A**). The six-rowed genotypes contributed to a higher share of genetic variation within populations as compared to two-rowed genotypes (**Table 2**). Six-rowed genotypes contained a higher number of polymorphic loci (Npl), of minor allele frequency (MAF), and polymorphic information content (PIC) than two-rowed genotypes. The allelic richness (Ar) and the number of private alleles were higher in two-rowed genotypes. However, equivalent values were observed between row types for Nei’s unbiased gene diversity (He) (**Table 3**).

**Table 2.** AMOVA for the studied barley genotypes based on row type and geographic origin.

| **Source of variation** | **Degree of freedom** | **Sum of square** | **Mean square** | **Variance component** | **Percentage of variation** |
| --- | --- | --- | --- | --- | --- |
| **Row type** |  |  |  |  |  |
| **Among Populations** | 1 | 811,123 | 811,123 | 1,498 | 10.6 |
| **Within Population** | 528 | 13,340,728 | 25,267 | 12,633 | 89.4 |
| **Total** | 1,059 | 14,151,851 | 13,363 | 14,132 | 100 |
| **2 Row** | 291 | 7,702,274 | 26,468 | — | — |
| **6 Row** | 237 | 6,352,044 | 26,802 | — | — |
| **Geographic origin** |  |  |  |  |  |
| **Among Populations** | 8 | 1,350,325 | 168,791 | 1722 | 12.3 |
| **Within Population** | 505 | 12,378,668 | 24,512 | 12,256 | 87.7 |
| **Total** | 1,027 | 13,728,993 | 13.368 | 13,978 | 100 |
| **Africa** | 11 | 263,938 | 23,994 | — | — |
| **Australia** | 22 | 472,595 | 21,482 | — | — |
| **Canada** | 18 | 383,147 | 21,286 | — | — |
| **Europe** | 47 | 945,066 | 20,108 | — | — |
| **ICARDA** | 287 | 7,418,540 | 25,849 | — | — |
| **India** | 39 | 876,475 | 22,474 | — | — |
| **Latin America** | 5 | 84,437 | 16,887 | — | — |
| **Landrace** | 16 | 304,167 | 19,010 | — | — |
| **USA** | 60 | 1,454,025 | 24,234 | — | — |

**Table 3.** Population genetics parameters for each population individualized per row type and geographic origin.

|  | N | Npl | Npa | Ar | He | MAF (%) | PIC |
| --- | --- | --- | --- | --- | --- | --- | --- |
| **Row Type** |  |  |  |  |  |  |  |
| **Two Row** | 292 | 38,954 | 24,108 | 1.99 | 0.33 | 0.25 | 0.33 |
| **Six Row** | 238 | 39,180 | 3,750 | 1.98 | 0.33 | 0.27 | 0.35 |
| **Geographic Origin** |  |  |  |  |  |  |  |
| **Africa** | 12 | 33,426 | 0 | 1.62 | 0.34 | 0.28 | 0.37 |
| **Australia** | 23 | 33,189 | 0 | 1.53 | 0.28 | 0.20 | 0.27 |
| **Canada** | 19 | 33,158 | 4 | 1.55 | 0.30 | 0.21 | 0.28 |
| **Europe** | 48 | 35,737 | 56 | 1.50 | 0.26 | 0.19 | 0.26 |
| **ICARDA** | 288 | 40,172 | 3,140 | 1.65 | 0.34 | 0.26 | 0.34 |
| **India** | 40 | 35,919 | 10 | 1.57 | 0.30 | 0.22 | 0.29 |
| **Landrace** | 17 | 34,702 | 64 | 1.60 | 0.32 | 0.22 | 0.31 |
| **Latin America** | 6 | 23,819 | 0 | 1.48 | 0.28 | 0.17 | 0.24 |
| **USA** | 61 | 38,160 | 6 | 1.60 | 0.31 | 0.23 | 0.31 |

N, number of genotypes. **Npl**, number of polymorphic loci. **Npa**, number of private alleles. **Ar**, allelic richness. **He**, Nei's unbiased gene diversity. **PIC**, polymorphism information content.

To better describe the pattern of genetic diversity across the geographic origins, the “Other” group was removed and nine subgroups (Africa, Australia, Canada, Europe, ICARDA, India, Landraces, Latin America, and USA) were considered for analysis. The AMOVA of this grouping explained up to 12.32% of the total genetic variation (**Table 2**). The genotypes from ICARDA, USA, and Africa groups contributed the most to the genetic variation as reflected by the mean squares. The genotypes from ICARDA contained the highest number of polymorphic loci and of private alleles. The lowest value of allelic richness was found in the Latin America population (1.4), probably due to the low number of entries in the group, while the ICARDA population showed the highest value (1.65). The Nei’s gene diversity varied from 0.26 (Europe) to 0.34 (ICARDA and Africa). The lowest MAF (0.17%) and PIC (0.24) were obtained for the Latin America population, while the Africa population showed the highest values with 0.28 and 0.37, respectively (**Table 3**). However, in pairwise differentiation among geographic groups, the highest Fst and therefore the highest difference was found for India vs Europe (Fst = 0.26) followed by Landrace vs Europe comparison (Fst = 0.25). Large differences were also found between, Australia vs Landrace (0.23) and Australia vs India (Fst=0.22) **(Table 4).** These results confirm the distinctiveness of European population from the other populations as reflected also by the PCA (**Figure 3B**). The ICARDA group showed generally lower pairwise Fst values, and therefore genetic distance as compared to all the other groups, despite the relatively higher Fst values shown with the European, Canadian and Australian groups **(Table 4)**.

**Table 4.** Population differentiation calculated using the Nei’s genetic distance (upper diagonal) and the Pairwise Fst (bellow diagonal) between populations defined according to their geographical origin.

|  | Africa | Australia | Canada | Europe | ICARDA | India | Landrace | Latin America | USA |
| --- | --- | --- | --- | --- | --- | --- | --- | --- | --- |
| **Africa** |  | 0.105 | 0.092 | 0.129 | 0.053 | 0.082 | 0.084 | 0.135 | 0.074 |
| **Australia** | 0.131 |  | 0.126 | 0.103 | 0.098 | 0.152 | 0.178 | 0.126 | 0.098 |
| **Canada** | 0.100 | 0.183 |  | 0.102 | 0.091 | 0.120 | 0.115 | 0.090 | 0.044 |
| **Europe** | 0.189 | 0.175 | 0.170 |  | 0.126 | 0.170 | 0.171 | 0.041 | 0.084 |
| **ICARDA** | 0.048 | 0.134 | 0.117 | 0.181 |  | 0.050 | 0.058 | 0.137 | 0.059 |
| **India** | 0.096 | 0.218 | 0.168 | 0.257 | 0.072 |  | 0.059 | 0.178 | 0.093 |
| **Landrace** | 0.076 | 0.231 | 0.146 | 0.247 | 0.066 | 0.071 |  | 0.178 | 0.093 |
| **Latin America** | 0.120 | 0.153 | 0.083 | 0.010 | 0.138 | 0.210 | 0.191 |  | 0.086 |
| **USA** | 0.082 | 0.144 | 0.052 | 0.140 | 0.086 | 0.138 | 0.121 | 0.078 |  |

### 3.3. Population Structure of the global barley panel

Population subgrouping using sNMF, cross-validation and the cross-entropy criterion approaches gave further information regarding the genotypic structure of the panel studied. The cross-entropy criterion did not exhibit a minimum value or a clear plateau and steadily decreased at higher k values (**Supplementary Figure 1**). However, substantial reductions of the criterion could be observed from 2 clusters to 8. Using 70% membership as a threshold, entries were assigned to different subpopulations at each K. Individuals whose highest ancestry coefficient is less than 70% were considered as admixed. Germplasm groups defined by sNMF with the number of ancestral populations corresponded to discrete clusters in the PCA space (**Figure 4**). At K=2, 51% of the genotypes were assigned to SubPop2.1 and 23% were assigned to SubPop2.2, while the remaining genotypes (26%) were admixed (**Figure 4C**). SubPop2,1 was mainly composed of the six-row genotypes (84%) and two-row genotypes (16%) mostly from ICARDA, while SubPop2.2 was composed of two-row genotypes from Europe, USA, Canada, Australia, ICARDA and Latin America origins. Subpop2.1 split into two groups at K=4; SubPop4.3 consisted of 25 (4.71%) two-rowed genotypes from ICARDA, and Subpop4.4 included six-row genotypes (4.15%) mainly from ICARDA, USA, India, and Canada. Similarly, Subpop2.2 split into two groups at K=4; SubPop4.1 comprised 23 (4.34%) genotypes mainly from ICARDA and Australia, and SubPop4.2 consisted of 56 (10.57%) genotypes from Europe, Canada, USA origins (**Figure 4D**; **Supplementary Table S5**).

**Figure 4.**
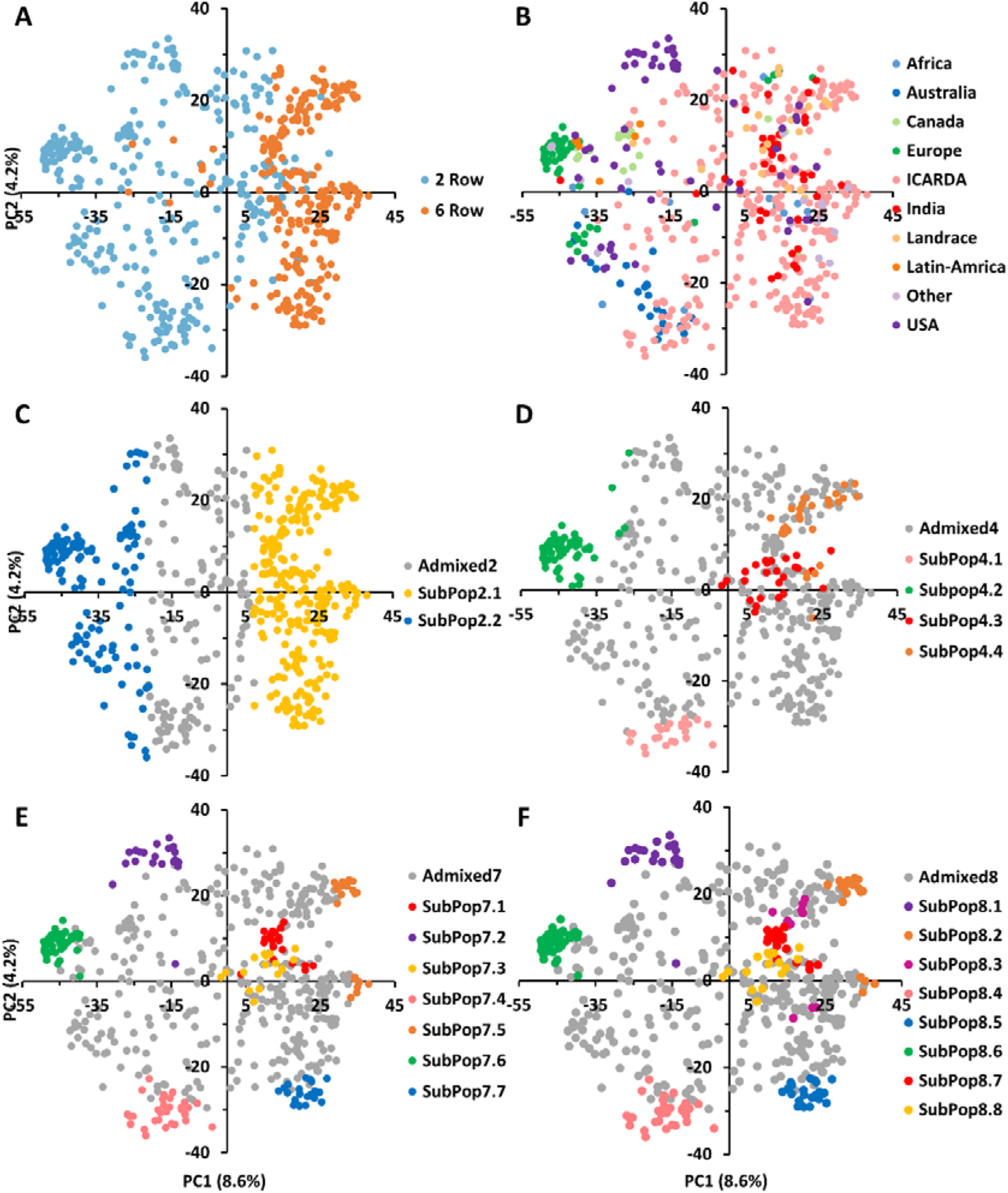
The genetic relationships and population structure of the 530 barley genotypes using SNP markers inferred by PCA and sNMF analyses. The same data points as in Figures 3A and 3B are shown and used for correspondence between population structure and PCA. Samples are colored according to their row type (A), geographic origin (B) and assignment to sNMF groups at K = 2 (C), K=4 (D), K=7 (E), and K=8 (F). genotypes whose highest ancestry coefficient is less than 70% are colored gray.

At K=7 (**Figure 4E**), two-rowed barleys located at the negative side of the first PC split into three subpopulations according to their origin; SubPop7.2 consisted of genotypes from USA, SubPop7.6 comprised genotypes from Europe, while genotypes from ICARDA and Australia were assigned to SubPop7.4, SubPop7.7 and Subpop7.5 located on the positive side of PC1 contained six-row genotypes from ICARDA while Subpop7.1 contained six-row genotypes from India and SubPop7.3 consisted of two-rowed genotypes from ICARDA. The population assignment at K=7 remained mostly constant at K=8 (**Figure 4F**). The new subpopulation (SubPop8.3) represented four genotypes from USA, four from Canada and one genotype from ICARDA. The SubPop8.4 harboring six-row genotypes from Canada (6), from ICARDA (12), India (4), USA (4) split into three subpopulations at K=8; SubPop8.2 representing ICARDA, SubPop8.3 representing Canada and USA and SubPop8.7 representing India. Over half of the genotypes placed in SubPop8.1 (SubPop7.5) were “PETUNIA 1” and its derived genotypes. Genotypes assigned to SubPop8.5 (SubPop7.7) were mostly issued from crosses with the cultivar “Rihane-03”. Similarly, genotypes included in SubPop8.8 (SubPop7.3 and SubPop4.3) were mainly derived from crosses with the cultivar “CANELA”. Most of the unassigned entries (admixed; n=356) observed in the collection were from ICARDA (58%) and to a lesser extent from USA (9.83%), and India (7.02%).

### 3.4. Genetic Diversity based on flowering loci

In addition to the genetic structure of the population, the diversity based on four SNPs previously identified as linked to major flowering genes and generating sixteen allelic combinations (AC) was studied. The AMOVA of the genetic differentiation between and within each flowering locus and their allelic combinations was conducted (**Table 5**). The marker associated to HvCEN explained the highest variation (11.11%) among populations followed by those associated to VRN-H3 (4.33%), while the VRN-H1 explained the lowest (2.53%) variation among populations. The sixteen allelic combinations explained 11.07% of total genetic variation. Fifteen genotypes with missing SNP data were not considered when calculating AMOVA based on allelic combinations. The sixteen allelic combinations were classified from the latest flowering (AC1) to the earliest (AC16; **Figure 5A**) on the base of the mean values of heading date recorded in 20 environments across Morocco, Lebanon, and India (data not shown).

**Figure 5.**
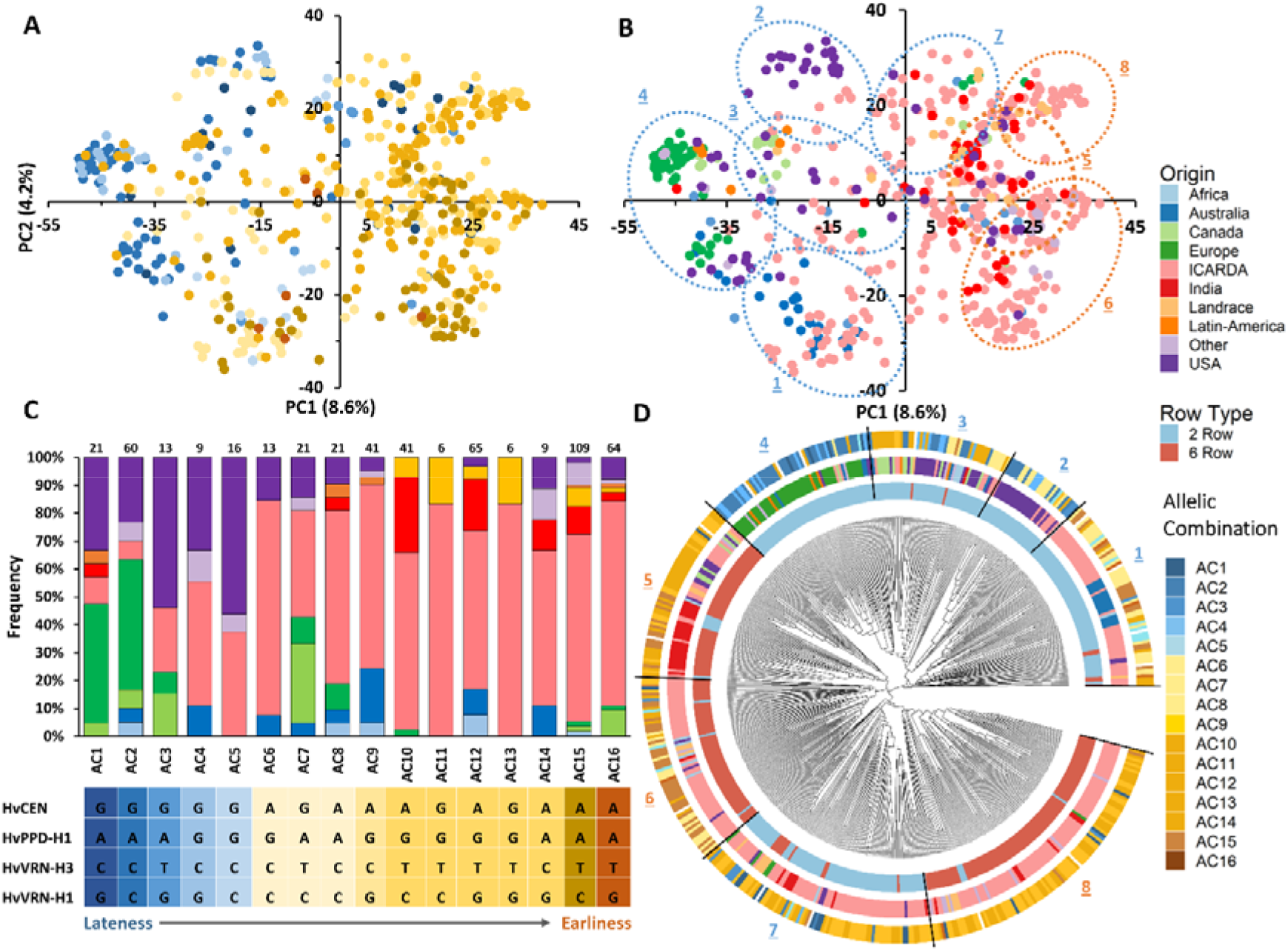
Principal component analysis (PCA) of the barley panel and distribution of allelic combinations (A) and geographic origins in the first two PCs (B). Histogram of allelic combinations frequencies in the nine geographical origins (C). NJ tree of the 530 barley genotypes based on 36,248 SNPs. The 530 accessions are colored according to their row type (inner circle) and geographic origin (medium circle), and allelic combination (outer circle).

All sixteen allelic combinations were present in the 275 genotypes from ICARDA origin. Thirteen allelic combinations were detected in USA group, eight in European, Australian and “Other” barleys, and seven in Indian genotypes. The most frequent allelic combinations in the germplasm were the early flowering AC15 (21.17%), AC12 (12.62%) and AC16 (12.43%). AC11 and AC13 were less frequent and unique to the ICARDA and Landrace populations. (**Figure 5C**; Supplementary Table S5). The PCA showed a clear distribution pattern of allelic combinations based on their effect on phenology. Genotypes with late allelic combinations were grouped on the negative side of the first PC while the early ones were more widespread (**Figure 5A**). Allelic combinations with late flowering characterized most two-rowed European, Canadian and USA genotypes and, to a lesser extent, the two-rowed African, Australian and Other origins (**Figure 5A, B**). The distribution of different allelic combinations within germplasm origin groups is shown in **Figure 5C**.

**Figure 5D** shows the assignment of allelic combinations to different clusters in the phylogenetic tree. The group of 2-rows genotypes comprises entries carrying late or intermediate flowering allelic combinations, mostly from USA, Canada and Europe origins belonging to groups 2,3 and 4, respectively. On the other hand, genotypes from ICARDA and Australia (cluster 1) carry early to intermediate flowering allelic combinations. Interestingly, Cluster 7, mainly composed of ICARDA entries, is characterized by early allelic combinations, with few genotypes having late flowering allelic combinations. The 6-row genotypes are mostly grouped in Clusters 5, 6 and 8 that are mostly composed of entries from ICARDA, India and other origins carrying early flowering allelic combinations. mostly USA, Canada and Europe origins respectively. In particular, the earliest allelic combinations (AC15 and AC16) were most frequent in ICARDA genotypes, while most European and USA barleys carried the latest flowering allelic combinations (AC1 and AC2). Two-rowed genotypes from USA carried mostly late flowering allelic combinations while in the six-rowed genotypes early flowering allelic combinations predominated. Frequent allelic combinations within Australian barleys were of the intermediate flowering AC9 (36%) and the early flowering AC12 (27%). Indian genotypes and Landrace included in this study carried predominantly the early flowering AC15 (28.2%; 43.8%), AC12 (10.8%; 18%) and AC10 (28.2%; 18%) respectively. These results indicate that the materials used in the analysis are carrying specific flowering allelic combinations linked to their geographic origins.

**Table 5.**
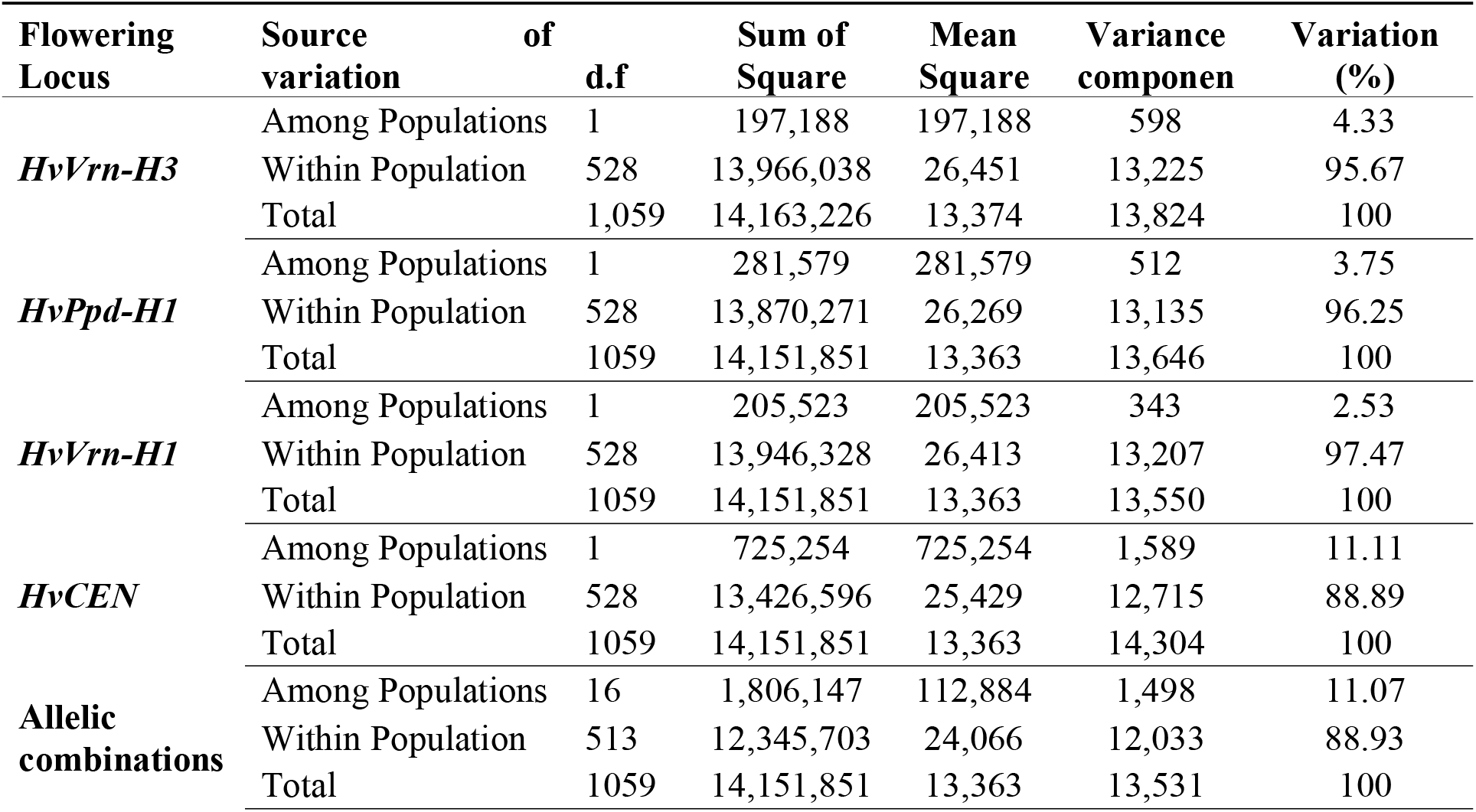
AMOVA of the 530 barley genotypes according to the allelic variants of main flowering genes.

### 3.5 Genetic sub-setting

To assemble a collection of diverse barley germplasm representative of the germplasm grown worldwide and with particular emphasis on Developing World a subset of the full collection was selected. The subsetting was done using the Mean of Transformed Kinships method aiming at a final number of 250 entries. The selection was made among the 312 entries of the panel that are of public domain and relevant to the developing world, mostly cultivars and breeding lines of ICARDA origin, both from the Syria and Latin America programs, and landraces (**Figure 6**). Up to 99.72% of the polymorphic markers present in the GBP set were also polymorphic in the new CGIAR subset. The distribution of the missing markers showed chromosome 7H as the one with less coverage (31) while chromosome 1H showed the highest coverage with only 3 markers missing. Moreover, 426 of the 471 rare alleles (alleles carried by <1% of the lines) of the total population are present in the subset. After filtering for minor allele frequency (MAF≥5%) and missing data (<10%), the number of polymorphic markers in the CGIAR set was 35,854, that is, 89% of the total number of polymorphic markers and 98.9% of the ones present after filtering in the GBP.

**Figure 6.**
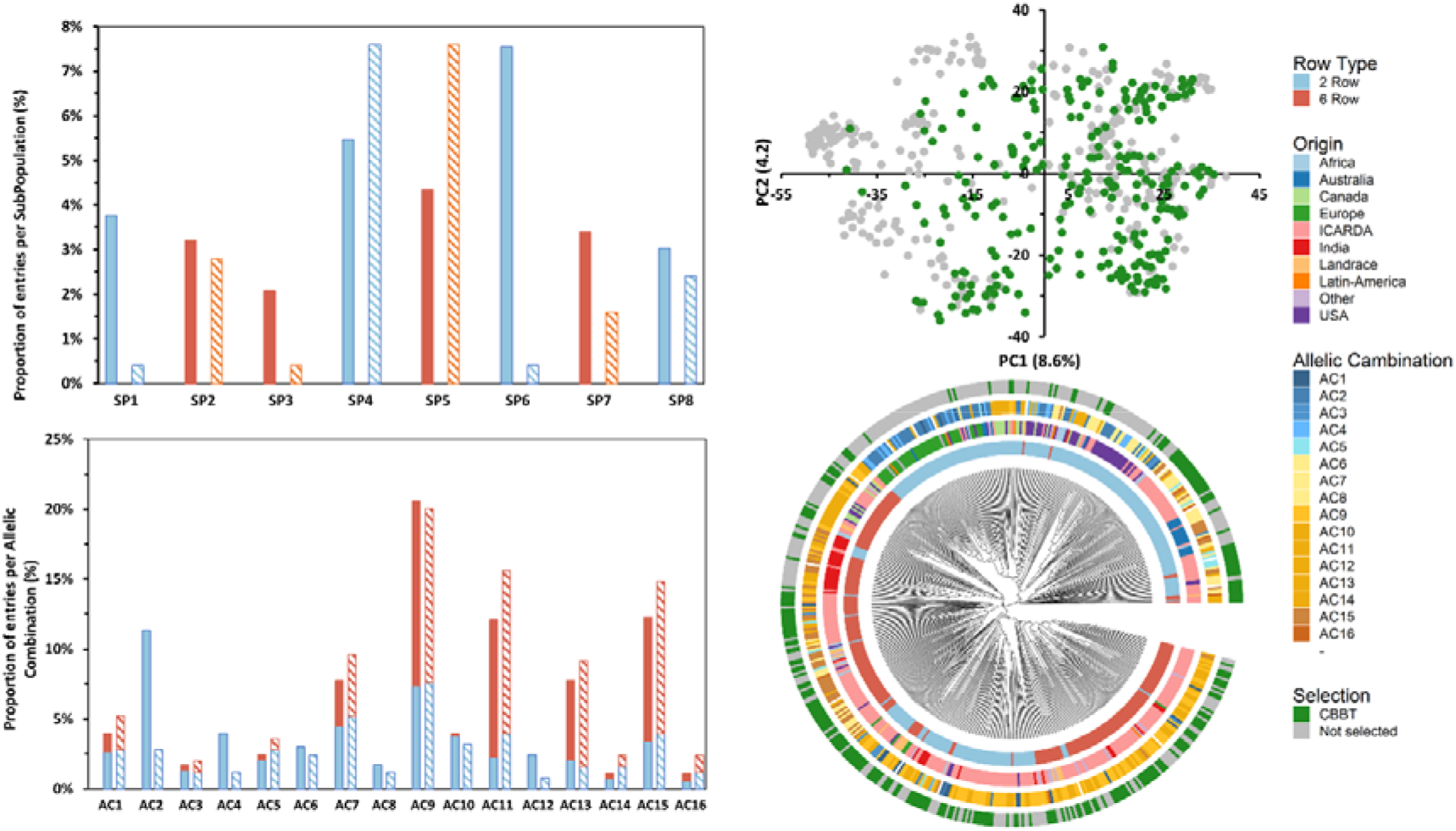
Principal component analysis (PCA) of the global barley panel and distribution of selected 250 entries of the CGIAR Barley Breeding Toolbox (CBBT) (B). Histogram of subpopulation (SP; A) and allelic combinations (AC; C) frequencies of the 250 entries of the GBP (bold bars) and the CBBT (striped bars) separating 6-row (orange) and 2-row (light blue) types. NJ tree of the 530 barley genotypes based on 36,248 SNPs. The 530 accessions are colored according to their row type (first and most inner circle), geographic origin (second circle), allelic combination (AC; third circle) and belonging to the CBBT set (fourth and most outer circle).

The subset continues to preserve a good balance between the row type with a slightly larger proportion of 6-row types (55% of the 250 entries) over the 2-rowed types as compared with the Global population (46% 6-row and 54% 2-row) (**Figure 6**). The selected subset showed a good coverage of most of the PCA spectra except the most negative values in PC1, where the European, Canadian and USA varieties are located (**Supplementary Table 6**). In fact, entries from all the SubPopulations identified in the present study are enclosed in the subset. The proportions are similar in the subset as compared to the Global Panel except for SubPopulation 1, 3 and 6 in which the subset shows a smaller proportion as compared with the GBP (**Figure 6**). These 3 SubPopulations are characterized by harboring entries from USA, Canada and Europe, the 3 origins identified as some of the least related to ICARDA material (Table 3). It is noteworthy to mention that all the Allelic Combinations identified at the most important phenological loci are represented in the subset (**Figure 6**).

## 4. Discussion

### 4.1. Genetic diversity

The characterization and dissection of genetic diversity and structure present in current germplasm collections help the breeders to meet breeding goals through the identification of beneficial alleles for traits of interest associated to molecular markers (Tester and Langridge, 2010; Al-Abdallat et al., 2017; Mazzucotelli et al., 2020). With this aim, in this study we explored the genetic diversity of a large global barley panel, comprising two- and six-rowed barley, from nine geographic origins and with different end-uses. The size and diversity of the panel used makes it a relevant tool for exploring the genetic diversity available for breeders.

Out of the 43,461 scorable SNPs markers of the 50k iSelect SNP array (Bayer et al., 2017), 40,342 SNPs were found polymorphic in the global barley panel. Thus, 92.5% of all potential markers were segregating in our panel and 89.8% of them did so in a significant number of genotypes (MAF>5%) and with minimum data loss (missing data<10%).

This result is similar and often higher than other reports of 39,733 SNPs (Darrier et al., 2019); 33,818 SNPs (Novakazi et al., 2020) and 37.242 SNPs (Cope et al., 2020). The highest number of SNPs was found on chromosomes 5H and 2H, respectively, while chromosome 1H contained the least number of SNPs accordingly with Bayer et al. (2017). Using the filtered set, we reported high gene diversity (0.35) and PIC (0.35), which indicates a similar level of genetic diversity as compared to previously studied collections. Higher PIC values have been reported for 170 Canadian cultivars and breeding lines (PIC=0.38; Zhang et al., 2009) and wild and cultivated barley (PIC= 0.38 and 0.35 respectively; Dai et al., 2012). Recent genomic studies reported lower gene diversity and PIC values in diverse germplasm collections including landraces, wild and cultivated barley (Pasam et al., 2012; Bengtsson et al., 2017; Amezrou et al., 2018; Verma et al., 2021). The high level of genetic diversity reported is probably a result of the wide range of worldwide germplasm collected across years and of the different objectives of the breeding programs. Particularly, the large proportion of ICARDA Global Barley Breeding Program elite lines contributed greatly to genetic diversity. This is probably due to its extensive use of genetic resources from the ICARDA Genebank and from the extensive breeding efforts and international collaborations and germplasm exchange needed to address the needs of a global breeding program.

### 4.2. Genetic diversity and population differentiation

Principal components analysis (PCA) and the NJ clustering partitioned the 530 entries into two clearly defined groups based on row-type (2-row vs 6-row) and highlighted the presence of subgroups characterized by common or closely related origins. Furthermore, AMOVA analysis indicated that germplasm origin could also explain a significant part of genetic diversity. The variation explained by geographic origin was higher than the value reported by Jilal et al. (2008) for a worldwide collection of 304 barley landraces from seven countries. The row type grouping captured 10.6% of the total genetic variance, which was lower than Malysheva-Otto et al, (2007) for 504 European barley cultivars released during the 20th century suggesting that the diversity of this panel goes beyond the row type genepools.

Row type in barley has been shown in the past to be one of the main determinants of the genetic structure, both due to the inherent nature of the trait which is associated to major regional preferences and its association to the different barley end-uses (Komatsuda et al., 2007; Saisho et al., 2009; Zhou et al., 2012). Most European countries exclusively use two-row barley for malt, while in North America, six-row varieties have played an important role in brewing (Verstegen et al., 2014; Milner et al., 2019). When comparing the genetic diversity, the six-row group showed more Npl, Nua, Ar, MAF, PIC values and genetic diversity relative to the number of genotypes as compared to the 2-row group. Higher MAF and PIC of six-row barley was also reported for a global barley panel (Hill et al., 2021), while the opposite was true for a Canadian barley collection (Zhang et al., 2009).

The group of ICARDA germplasm showed high genetic diversity as revealed by the number of polymorphic loci, number of unique alleles and Nei’s unbiased gene diversity. Moreover, 99.72% of the 40,342 polymorphic SNPs in the panel were so in the ICARDA population too. Interestingly, the lower differentiation between ICARDA germplasm and most of the other origins evidenced the long-lasting collaborations between the Global Program and national and regional breeding programs. In fact, more than 250 spring and winter 2-row, 6-row and naked barley varieties of ICARDA origin have been released in 46 countries since 1979, including USA (5 varieties), Canada (15 varieties), Australia (4), India (8 varieties) and some African (70) and Latin American (27) countries among others. ICARDA germplasm has in addition being extensively used in crosses in the National programs. Interestingly, the Indian germplasm is closely related to the groups of Landrace, ICARDA and Africa, while distinct from the other groups. Landrace group exhibited the lowest Npl, MAF and PIC values which may be explained by the under-representation of Landrace genotypes during the development of the 50K SNP array (Bayer et al., 2017; Darrier et al., 2019).

Latin American germplasm exhibited the highest He, MAF, PIC although having the lowest number of individuals (5) as compared to other origins. This could be due to the diverse source of genotypes originating from different countries (Argentina, Peru, Ecuador and Uruguay). Despite American malting varieties being obtained through hybridization with European accessions, the national industrial standards for malting and brewing have led to the development of germplasm with specific characteristics (Matus and Hayes, 2002). The European barley germplasm showed the lowest allelic richness and genetic diversity as compared to other geographical origins. Similar findings were achieved in a previous study by Malysheva-Otto et al. (2006) when conducting a genetic diversity study with 953 barley accessions and 48 SSR markers. Furthermore, it has been suggested that the replacement in Europe of six-row spring barley by two-row spring barley during the 18th and 19th centuries led to a loss of diversity (Backes et al., 2003). In addition, the European varieties used were also the least related to the ICARDA germplasm. This could be due to the type of environments these varieties target in Europe. Most of the European varieties used in the present study were released in West, Central and Northern Europe where the type of environment and stresses associated differ from the main target areas of the Global Barley Breeding Program. In fact, while there have been collaborations between ICARDA and research groups from these countries both nowadays and in the past, practically all 25 ICARDA originated varieties released in Europe have been so in countries of the Mediterranean Basin.

### 4.3. Flowering genes drive the geographic adaptation of barley

In the current study, we also analyzed the allelic diversity at four SNPs associated with key flowering genes (*HvCEN, HvVrn-H3, HvPpd* and *HvVrn-H1*) and their additive effect on the phenology and adaptation of the panel studied. These highly polymorphic (PIC > 0.4) loci were not in LD (**Supplementary Table S2**) and showed to be associated to the genetic structure of the population. The different allelic combinations of the four markers used explained 11.07% of the total variance and the distribution of *HvCEN* was the main driver of the association between the flowering genes and genetic structure of the population with minor effects of the other flowering genes studied. This influence was even more evident in the PCA graph which, in addition, shows clear geographical structuring based on earliness/lateness of the constructed allelic combinations. Comadran et al. (2012) and Russell et al. (2016) found that the patterns of single and multiple-flowering gene combinations contributed to a wide range of ecogeographical adaptation in barley.

The ICARDA group contained all sixteen allelic combinations indicating high allelic variability that could provide adaptation to different environments and farming practices. Notably, in the rest of the germplasm, most of the genotypes harboring early-flowering allelic combinations are six and two-row barleys from India, ICARDA and Australia, while most of the accessions with late allelic combinations grades are two-rowed barleys from Europe, USA, Canada and Latin America. However late-flowering allelic combinations were also present in low frequencies in two-rowed genotypes from Africa, Australia, and ICARDA.

Alqudah et al. (2014) found a similar separation of a spring barley collection based on the response to photoperiod and reduced photoperiod sensitivity (Ppd-H1/ppd-H1) at heading time. In this study, most of the accessions in the PpdH1-group were six-rowed barleys from West Asia and North Africa (WANA) region and East Asia (EA), while most of the accessions in the ppd-H1 group were two-rowed barleys from Europe. Spring barley accessions originating from regions where the growing season is short and with a dry summer (WANA, India e.g.) tend to carry photoperiod responsive Ppd-H1 alleles, causing early heading under LD, while late-flowering allelic combinations in European and North American accessions of spring barley could be due to reduced photoperiod sensitivity, ppd-H1, alleles supporting the results of (Turner et al., 2005; Wang et al., 2010; Alqudah et al., 2015).

Saisho et al. (2011) highlighted the geographic distribution of vernalization requirements in domesticated barley; western regions including Turkey, Europe, North Africa, and Ethiopia were strongly associated with a higher degree of spring growth habits, while the extreme winter growth habits were localized to Far Eastern regions including China, Korea and Japan. In the European-cultivated germplasm, most variation in vernalization requirement is accounted for alleles at the VRN-H1 and VRN-H2 loci, as the majority of European varieties are thought to be fixed for winter alleles at the VRN-H3 locus (Cockram et al., 2007). In the Landrace group predominated early-flowering allelic combinations and showed an allelic profile similar to India and six-rowed genotypes representing CWANA region. The barley landraces have adapted to seasonal photoperiod and temperature by changing their growth habit from a facultative winter-planted species in its center of origin to a short-season spring-planted crop on the northwestern fringes of Europe and highland plateaus of Central Asia (Comadran et al., 2012).

### 4.4. Population structure

It is generally agreed that the principal sources of structure in diverse collections of barley germplasm are spike row number, growth habit, and geographic origin (Cuesta-Marcos et al., 2010; Hamblin et al., 2010; Comadran et al., 2012). The results obtained through population structure, principal component analysis, neighbor-joining clustering analysis indicate a separation based on inflorescence type, geographic origin and phenology. Generally, PCA does not always classify accessions into discrete clusters, especially not when admixed accessions and accessions of various geographical origins are included (Glaszmann et al., 2010). In our case, subpopulations defined by sNMF correspond to discrete clusters in the PCA space. At K=2, a division according to row type was the main driver. Finer subdivisions at higher K could be interpreted based on geographical origin, phenology and row type generally within the discrete clusters established by the phylogenetic tree. Earlier studies reported similar population divisions (Muñoz-Amatriaín et al., 2014; Al-Abdallat et al., 2017; Bengtsson et al., 2017; Amezrou et al., 2018; Milner et al., 2019; Verma et al., 2021).

Two-row cultivars from Australia, USA and Canada alongside two-rowed varieties from Latin America and Europe were mostly grouped in the first, second, third and the fourth clusters, respectively. This relatedness can be explained by the European origin of American cultivated barley that was introduced in the continent from Europe nearly 500 years ago. Then after, the similar climate and continual human migration has likely led to subsequent introductions and exchanges (Casao et al., 2011). This relatedness could be seen already since the K=2 separation population structure. However, at higher Ks, the USA and Australian, most of them malting barley types, formed relatively distinct and stable groups within the diversity observed across the panel and had little overlap with the genotypes from other groups. This was especially true also for the European two-rowed barleys. Malysheva-Otto et al. (2006) suggested that the lower genetic diversity existing in European barley may be explained by the fact that exotic varieties were very rarely involved in the breeding programs in Europe. It is also important to note that two-rowed barleys are mostly malting types which may lead to specialized breeding gene pools. Two-rowed genotypes grouped in cluster 7 were from ICARDA origin and showed also distinctiveness at least from K=4 (SubPop4.3). These lines included the cultivar *CANELA* and its derived crosses (**Supplementary Table S4**). *CANELA* is a 2-row malt barley resistant to at least 7 diseases, issued from the ICARDA/CIMMYT Latin America breeding program where the focus on malt barley was particularly strong as compared to the Syrian program.

Cluster 5, 6 and 8 assembled most of the 6-row barley lines of the panel. Although in this case, the phylogenetic clustering did not completely overlap with the subpopulations at K=4, at larger K these three clusters were evident. Cluster 5 grouped two subpopulations at K=8. The first subpopulation consisted of USA and Canada 6-row feed barley entries. The second grouped mostly 6-row Indian varieties. Cluster 6 instead consisted of a distinct subpopulation from ICARDA origin (SubPop7.7 and SubPop8.5). Genotypes of this group were mostly derived from crosses with the cultivar *RIHANE-03*, a mega-variety from the ICARDA Syria program released in several North African and West Asian countries characterized by its drought tolerance. Cluster 8 harbored one subpopulation (SubPop7.5 and SubPop8.2). This subpopulation assembled almost exclusively the 6-row ICARDA lines issued from the ICARDA/CIMMYT Latin America program, notably the lines derived from the cultivars *PETUNIA-1* and, especially, *‘DOÑA JOSEFA’* (aka *V-MORALES*) a widely adapted 6-row malt barley line. The last subpopulation consisted of genotypes mostly issued from crosses between entries from the Syrian and Mexican programs. Despite this differentiation, the PCA showed some level of overlap in Clusters 5, 6 and 8 suggesting that these subpopulations may have some genetic overlapping that goes beyond the origin.

Admixed genotypes at K=8 belonged to nine geographical origins and 57.6% of them were ICARDA breeding lines (47% two-row and 53% six-row). The higher number of admixed genotypes among the ICARDA lines can be explained by their largest population size (53.6% of genotypes present in the collection) and its extensive use of two-by-six row crosses for germplasm improvement (Amezrou et al., 2018; Visioni et al., 2018; Verma et al., 2021). Likewise, two-by-six row crosses hybridization is routinely practiced in India for improving the adaptability of exotic two-row barleys with indigenous six-row cultivars (Verma and Sarkar, 2010).

### 4.5. CGIAR Barley Breeding Toolbox

The main objective of the present study was to identify and assemble an Association Mapping panel of barley genotypes that represented the diversity of the Global Barley Panel to serve as a CGIAR barley breeding toolbox (CBBT) especially for the Developing World. For it, the main criteria to select among the lines in the Global panel was: i) to be representative of the germplasm grown in the Developing World; ii) to cover a wide range of genetic variability, morphophysiological relevant traits and phenology groups and iii) be of public domain or international public goods to facilitate germplasm exchange.

In order to meet these criteria and due to their large diversity, the CGIAR entries, including landraces hosted in the ICARDA genebank were pre-selected. The Developing World representativity requirement was met due to the global target of the ICARDA Global Barley Breeding program of the CGIAR especially in terms of adaptation and end-uses. In fact, most of the CGIAR lines included in the panel have been shared with and selected by National Partners as part of the Barley International Nurseries. These sets of diverse and elite barley genotypes distributed upon request to more than 70 collaborators in 20 countries in America, Europe, Africa and Asia (Sanchez-Garcia et al., 2021) annually have resulted in the release of more than 250 varieties since 1977, mostly in the Developing World. Moreover, these genotypes are transferred as International Public Goods under a Standard Material Transfer Agreement that allows their free use for research, education, and breeding. The CGIAR lines also showed wide diversity and representativity of the whole population. This group of lines showed the largest number of polymorphic loci and together with the landraces covered most of the diversity in the population. Moreover, the CGIAR origin lines are present in all the subpopulations and showed all 16 allelic combinations of phenology genes.

However, to facilitate future phenotyping studies and reduce redundancies a core subset of 250 entries was selected using Mean of Transformed Kinships method. The use of this method differs from others like the PIC based one used in other studies such as Muñoz-Amatriaín et al. (2014) due to the different aim of the subset. In the case of the CBBT, the primary aim is to serve as an Association Mapping panel for researchers that do not have access to one and rare trait discovery is only a secondary objective. However, the subset chosen kept most of the diversity of the Global panel and its morphophysiological traits’ proportions suggesting that rare trait discovery is still possible. In summary, the CBBT is a solid and diverse representation of the barley grown in developing countries but also significant in the developed World.

The genotypic data of the population is available in Supplementary Table 7 and in *Germinate* Database (https://germinateplatform.github.io/get-germinate/). All the data generated on this population will be freely available in Germinate for the global barley community to use.

## 5. Conclusion

The main objective of the present study was to assemble a collection of diverse barley germplasm representative of the germplasm grown worldwide and with particular emphasis on Developing World. The results showed that the Global panel used in this study is highly diverse and covers a wide range of both genetic variability of cultivated barley and phenology groups. The CBBT assembled represents with fidelity the genetic diversity and key morphophysiological traits of the Global panel. This collection will be made available, together with the genotypic data, to breeders and researchers worldwide to serve as a collaborative tool to underpin the genetic mechanisms of traits of interest for barley cultivation. In addition, the collaborative approach to gene discovery will make this collection a toolbox for breeders that will be able to use the germplasm and the markers identified by researchers worldwide in their own breeding programs.

## 6. Acknowledgments

The present work was supported by the *Modernization of crop breeding programs at ICARDA - building on past AFESD support to prepare for the future* project funded by the Arab Fund for Economic and Social Development (AFESD). The authors would like to thank the USDA small grains genotyping lab in Fargo (North Dakota, USA) and Dr Shiaoman Chao for the genotyping of 266 genotypes used in the present study. The authors acknowledge the support of the CGIAR SAPLING and ABI initiatives.

