## Supplementary figures and images for "CGIAR BARLEY BREEDING TOOLBOX: A diversity panel to facilitate breeding and genomic research in the Developing World"

### Supplementary Figure 1

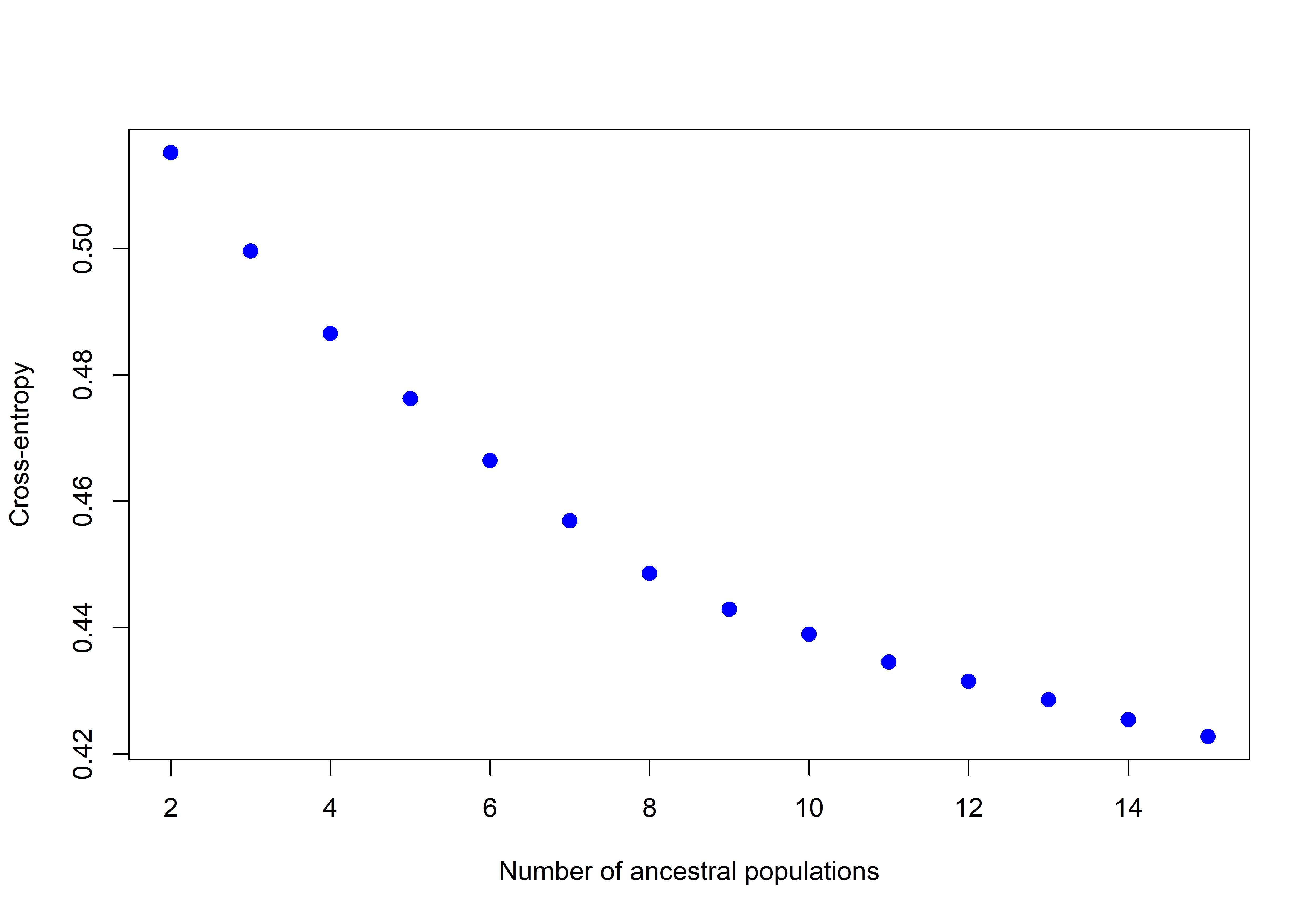
